# Everyday taxi drivers: Do better navigators have larger hippocampi?

**DOI:** 10.1101/431155

**Authors:** Steven M. Weisberg, Nora S. Newcombe, Anjan Chatterjee

## Abstract

Work with non-human animals and human navigation experts (London taxi drivers) suggests that the size of the hippocampus, particularly the right posterior hippocampus in humans, relates to navigation expertise. Similar observations, sometimes implicating other sections of the hippocampus, have been made for aging populations and for people with neurodegenerative diseases that affect the hippocampus. These data support the hypothesis that hippocampal volume relates to navigation ability. However, the support for this hypothesis is mixed in healthy, young adults, who range widely in their navigation ability. Here, we administered a naturalistic navigation task that measures cognitive map accuracy to a sample of 90 healthy, young adults who also had MRI scans. Using a sequential analysis design with a registered analysis plan, we did not find that navigation ability related to hippocampal volume (total, right only, right posterior only). We conclude that navigation ability in a typical population does not correlate with variations in hippocampal size, and consider possible explanations for this null result.

## 1. Introduction

Spatial navigation is a fundamental problem faced by any mobile organism. This ability is supported in part by the hippocampus, which is theorized to construct a cognitive map – knowledge of the distances and directions between landmarks (O’Keefe & Nadel, 1978; Tolman, 1948). Evidence for the role of the hippocampus in navigation comes from functional neural data across a wide range of levels of analysis. At the single-cell level, place cells in the hippocampus fire when an animal is in a certain location (Ekstrom et al., 2003; O’Keefe & Nadel, 1978). At the voxel level, fMRI reveals that the patterns of voxels in the hippocampus map to information about spatial distance (Vass & Epstein, 2013). At the whole hippocampus anatomic level, fMRI reveals that the hippocampus is more active during active navigation than passive travel or when following a familiar route (e.g., Hartley et al. 2003). So, neuronal activity in the hippocampus is consistent with the hypothesis that the hippocampus constructs a cognitive map.

Intriguingly, there is reason to suppose that variations in the structure and function of hippocampus may relate to variations in navigation abilities. Maguire and colleagues (2000; 2006) showed that the right posterior hippocampus was enlarged in taxi drivers from London, who memorize an enormous catalog of spatial information and navigate easily around the complex layout of London. Further work showed that elderly taxi drivers who were still driving taxis had enlarged right posterior hippocampi compared to elderly taxi drivers who had stopped (Woollett, Spiers, & Maguire, 2009). Although this work on taxi drivers presented among the first data to show a correlation between hippocampal volume and aspects of navigation behavior, the sample sizes were small (less than 20 participants in most studies). And despite making claims about increased right posterior hippocampal volume and decreased right anterior hippocampal volume in taxi drivers, the researchers did not test the key interaction between posterior and anterior hippocampal volume between taxi drivers and controls, rendering this conclusion unsupported by the data. Nevertheless, complementary research in non-human animals has supported the notion that larger hippocampi are associated with better navigation (Sherry, Jacobs, & Gaulin, 1992). Importantly, changes in hippocampal size may occur within an animal’s lifespan. Male meadow voles, for example, show cell proliferation in the dentate gyrus concomitant with hippocampal volume increases in the breeding season (when males have greater spatial navigation requirements) compared to the non-breeding season (Galea & McEwen, 1999). Similarly, compromise of the hippocampus is associated with spatial navigation deficits. Poor spatial navigation occurs in patients with Alzheimer’s disease (Deipolyi, Rankin, Mucke, Miller, & Gorno-Tempini, 2007; Konishi et al., 2018; Moodley et al., 2015; Plancher, Tirard, Gyselinck, Nicolas, & Piolino, 2012), patients with brain lesions (Kolarik, Baer, Shahlaie, Yonelinas, & Ekstrom, 2018; Rosenbaum et al., 2000; Smith & Milner, 1981) and in the elderly (Konishi, Mckenzie, Etcharnendy, Roy, & Bohbot, 2017; Moodley et al., 2015). These groups show reduced hippocampal volume, suggesting that a healthy hippocampus is critical for normal spatial navigation function.

Collectively, this body of literature seems to make a powerful case for an association between hippocampal volume and navigation. However, evidence from healthy young adults is more mixed. Some research studies are positive. First, there are findings that hippocampal volume correlates with specific navigationally-relevant spatial tasks, notably perspective taking. (in which an unseen view must be imagined). Perspective taking correlates with spatial memory for large-scale environments (Weisberg & Newcombe, 2016) and elicits neural activation in the hippocampus (Lambrey, Doeller, Berthoz, & Burgess, 2012). Hartley and Harlow (2012) created a perspective-taking task in which participants match an image of three-dimensional mountains to an image of the same mountains range viewed from a different perspective while ignoring similar-looking foils. Accuracy on this task correlated with bilateral hippocampal volume. Sherrill and colleagues (2018) measured participant’s ability to find a goal from first-person and map perspectives after viewing a map with their position and the position of the goal. Accuracy in the first-person condition correlated with bilateral posterior hippocampal volume. Second, hippocampal volume correlates with specific strategies used by healthy young adults when navigating. Bohbot and colleagues measured spatial navigation strategy using a task in which the direction to goals could be based on which response should be made (e.g., to one’s right; thought to rely on the caudate nucleus), or based on each goal’s position relative to external landmarks (e.g., the church is to the right of the school; thought to rely on the hippocampus), independent of success at finding a goal. They found large positive correlations with hippocampal volume and the number of goals found relative to external landmarks (Bohbot, Lerch, Thorndycraft, Iaria, & Zijdenbos, 2007). Third, using a self-report measure of navigation ability, two studies with large samples reported correlations with hippocampal volume, although the effect sizes were modest (Hao et al., 2016; Wegman et al., 2014).

None of these approaches is direct, however. Perspective taking is only one component of successful navigation. Navigation strategy can be orthogonal to navigation accuracy (Marchette, Bakker, & Shelton, 2011). Self-report is correlated with, but not the same as, navigation accuracy. In a more direct look at the issue in typical adults using a real-world environment to measure navigation ability found a large correlation with right posterior hippocampus. However, that study suffers from the use of a small sample (Schinazi, Nardi, Newcombe, Shipley, & Epstein, 2013), and we are unaware of other studies of this kind. Additionally, studies in this literature vary in how they define the relevant hippocampal areas, analyzing right posterior hippocampal volume (Maguire et al., 2000, 2006; Schinazi et al., 2013), total hippocampal volume (Hao et al., 2016; Hartley & Harlow, 2012; Konishi et al., 2017; Wegman et al., 2014) or both posterior hippocampi (Sherrill et al., 2018). Moreover, some studies correct for total brain volume, gender, and age, whereas others do not. These different anatomic and analytic choices undermine confidence in the premise that hippocampal structure correlate with navigation.

Here, we test the hypothesis that hippocampal volume is a biological marker for spatial navigation ability in young, healthy human subjects. We test this hypothesis in a large sample using a widely-used desktop virtual environment (Virtual Silcton; Weisberg et al. 2014; Weisberg and Newcombe 2016). Virtual Silcton measures navigational accuracy while allowing participants to vary in the strategy they use. As a primary behavioral measure of interest, we chose total pointing performance – or how accurately participants could point to and from all locations in Virtual Silcton. This measure captures the accuracy with which participants learned the direction from each building to every other, and may substitute for the ability to take a novel shortcut – a hallmark of the cognitive map. Unlike the task used by Bohbot and colleagues (2007), the pointing task used in Virtual Silcton does not constrain participants to use one navigation strategy or another.

We chose right total hippocampal volume as the primary target of analysis because right hippocampal volume is more consistently reported to be related to navigation ability than left. We thus chose right hippocampal volume as our primary confirmatory analysis, the simplest measure of hippocampal volume, which would not introduce additional issues of reliability in segmentation or choice of measurement technique. We registered one confirmatory analysis using a sequential analysis design (Lakens, 2014) in which we planned to correlate right total hippocampal volume with how well participants learned locations after navigating in Virtual Silcton. However, whereas some data suggest that the posterior hippocampus on the right relates most strongly to spatial navigation ability (Maguire et al., 2000; Schinazi et al., 2013), other research shows this relationship with the right anterior hippocampus (Wegman et al., 2014) or right total hippocampus (Hao et al., 2016; Hartley & Harlow, 2012; Konishi et al., 2017). Some research has assessed as the primary measure of interest the ratio between anterior and posterior hippocampus (Poppenk, Evensmoen, Moscovitch, & Nadel, 2013). For that reason, in exploratory analyses, we also looked at anterior, posterior, and total hippocampal volume on the right and left.

We also considered the possibility that non-hippocampal brain structures might relate to navigation ability, or that alternative measures of navigation ability might better capture hippocampal-based navigation. We thus conducted exploratory analyses relating accuracy on several Virtual Silcton measures (subsets of the pointing task, a map constructed from memory, and building naming) and non-Silcton measures (including mental rotation, verbal ability, self-reported navigation ability, and self-reported spatial anxiety) to the volume of various brain structures (including sub-divisions of left and right hippocampi, caudate nucleus, the amygdala, and total cortical volume).

## 2. Materials and Methods

### 2.1. Participants

We recruited participants by advertising to and recruiting from participants who had participated in fMRI experiments from the Center for Cognitive Neuroscience at the University of Pennsylvania, asking them to participate in a one-hour study for which they would be paid $10.

We recruited 90 participants (54 women). Nineteen participants self-reported as Asian, 17 as African-American or Black, and 42 as Caucasian or White. Thirteen participants self-reported as Hispanic, three reported multiple races, one reported other, and one participant did not report ethnicity or race. Participants’ average age was 23.1 years (*SD* = 3.94).

### 2.2. MRI Acquisition

Scanning was performed at the Hospital of the University of Pennsylvania using a 3T Siemens Trio scanner equipped with a 64-channel head coil. High-resolution T1-weighted images were acquired using a three-dimensional magnetization-prepared rapid acquisition gradient echo pulse sequence. Because these data were collected for different research studies, specific parameters varied by protocol (see Supplementary Table 1).

### 2.3. Volumetry Measures

We calculated neuroanatomical volume of cortical structures in two ways. For the main analysis of the right hippocampus, we extracted hippocampal volume in two ways – Freesurfer and Automatic Segmentation of Hippocampal Subfields (ASHS). For the exploratory analyses, including sub-regions of the hippocampus and additional neuroanatomical structures, we focus on the parcellation from ASHS in the main text, but include analyses from Freesurfer in the supplementary results.

We used Freesurfer 6.0 (Iglesias et al., 2015) software to extract volume estimates of cortical and subcortical regions as part of the standard recon-all pipeline. We segmented posterior and anterior hippocampus manually using Freesurfer’s hippocampal parcellation. Anterior hippocampus was defined as all voxels in this parcellation that were in all slices anterior to (and including) the last coronal slice with at least 3 pixels that could be identified as the uncus (as defined in Morey et al. 2009). We then segmented posterior and anterior hippocampus manually using Freesurfer’s hippocampal parcellation. Anterior hippocampus was defined as all voxels in this parcellation that were in all slices anterior to (and including) the last coronal slice with at least 3 pixels that could be identified as the uncus (as defined in Morey et al. 2009). We also used the ASHS pipeline, which performs automatic parcellation of the hippocampus and other medial temporal lobe structures, including estimates of posterior and anterior hippocampus (Yushkevich et al., 2015).

### 2.4. Behavioral and Self-report Measures

#### 2.4.1. Demographics

We asked participants to report their biological sex, gender, ethnicity, age, education level, and handedness.

#### 2.4.2. Wide range achievement test 4 – verbal (WRAT-4; Wilkinson and Robertson 2006)

The WRAT-4 Word Reading Subtest is a measure of verbal IQ that correlates highly with the WAIS-III, and WISC-IV (Strauss, 2006). The WRAT-4 Word Reading Subtest requires participants to pronounce fifty-five individual words. Each participant’s score is the number of words pronounced correctly out of 55. Any participants who reported speaking any language besides English as their first language were excluded from these analyses (eight participants were excluded based on this criterion).

#### 2.4.3. Spatial anxiety questionnaire (SAQ; Lawton 1994)

This self-report measure of spatial anxiety consists of eight 7-point Likert-scale items that ask participants to indicate their level of anxiety when confronting situations such as “Locating your car in a very large parking garage or parking lot,” and “Finding your way to an appointment in an area of a city or town with which you are not familiar.”

#### 2.4.4. Santa Barbara sense of direction scale (SBSOD; Hegarty, Richardson, Montello, Lovelace, & Subbiah, 2002)

This self-report measure of navigation ability consists of fifteen 7-point Likert-scale items such as “I am very good at giving directions,” and “I very easily get lost in a new city.” The average score for each participant has been shown to correlate highly with performance on behavioral navigation tasks in real and virtual environments (Hegarty et al., 2002; Weisberg et al., 2014).

#### 2.4.5. Mental rotation test (MRT; Vandenberg and Kuse 1978; adapted by Peters et al. 1995)

This computerized version of the MRT consists of two 10-item sections of multiple-choice questions. Participants have three minutes per section. Each item consists of one target two-dimensional image of a three-dimensional shape made up of connected cubes, and four answer choices, also made up of connected cubes. Two of the answer choices are the same configuration of cubes, but rotated in 3D space. The other two answer choices are a different configuration. Participants received two points per correct choice, and lost two points per incorrect choice. Zero points were awarded for each omission.

#### 2.4.6. Virtual Silcton (Schinazi et al., 2013; Weisberg & Newcombe, 2016; Weisberg et al., 2014)

Virtual Silcton is a behavioral navigation paradigm administered via desktop computer, mouse, and keyboard. Modeled after the route integration paradigm (e.g., Hanley and Levine 1983; Holding and Holding 1989; Ishikawa and Montello 2006; Schinazi et al. 2013), participants learn two routes in separate areas of the same virtual environment by virtually traveling along a road indicated by arrows (see Figure 1). They learn the names and locations of four buildings along each of these routes. Then, they travel along two routes which connect the two areas from the first two routes. Virtual travel consisted of pressing arrow keys (or the W, A, S, and D keys) on a standard keyboard to move in the environment, and moving the mouse to look around. Participants were constrained to travel only along routes indicated with arrows. That is, we surrounded each route with invisible walls that restricted movement off the routes, but could be seen through. Participants could move and look at whatever pace they chose. Participants had the opportunity to learn each route once. At a minimum, we required participants to travel from the beginning to the end and back to the beginning of each route, but participants could spend as much time and do as much backtracking as they liked. Buildings were indicated by blue gems, which hovered over the path, and named with signs in front of the building.

**Figure 1.**
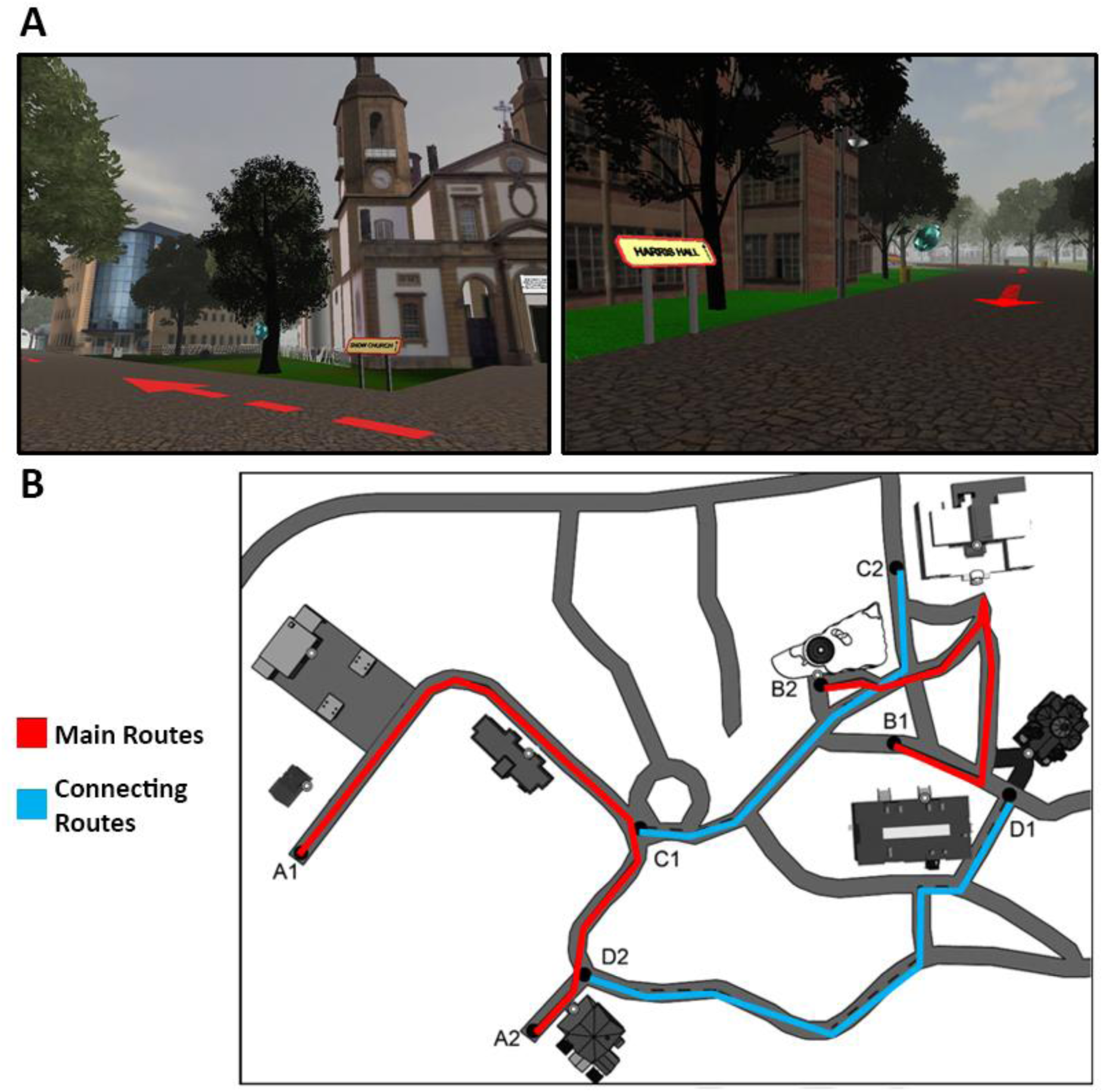
Screenshots and map of Virtual Silcton. Screenshots from Route A and B (A) and aerial map of Virtual Silcton, never seen by participants (B). Buildings were indicated by blue gems, which hovered along the path and named with yellow and red signs. Small white circles on the map indicate the front door of each building, which was the exact spot participants were asked to point to during the pointing task.

Participants were tested on how well they learned directions among the buildings within each of the main routes, and among buildings between the main routes. Testing involved two tasks. For an onsite pointing task, participants pointed to all buildings from each building they learned. The participant viewed the virtual environment along the route, next to one of the buildings they learned, and moved the mouse to rotate the view and position a crosshair toward one of the other buildings, then clicked to record the direction. The name of the building at the top of the screen then changed, and the participant pointed to the next named building. The dependent variable was calculated as the absolute error between the participant’s pointing judgment and the actual direction of the building (if this difference was greater than 180°, it was corrected to measure the shorter of the two possible arcs). We calculated pointing error separately for within-route trials and between-route trials separately. This resulted in 32 between-route trials and 24 within-route trials. Of the 24 within-route trials, 14 were mutually-intervisible (i.e., if any part of the building being pointed at was visible from the building being pointed from, it counted as a mutually-intervisible trial), while 10 were not.

Participants also completed a model-building task wherein they viewed a rectangular box on a computer screen and birds-eye view images of the eight buildings. Scrolling over the buildings with the mouse revealed a picture of the front view of the building and its name. Participants were instructed to drag and drop buildings to the position in the box they believed the building would be located (as if they were creating a map), without regard to the orientation of the buildings or to the map. The model-building task was scored using bidimensional regression analyses (Friedman and Kohler 2003).

Finally, in the building naming task, participants were shown pictures of each building and asked to name the building to the best of their ability.

#### 2.4.7. Debriefing and strategy questionnaire

We asked participants two debriefing questions: “What was the hardest part of the navigation test?” and “Did you have trouble remembering the names of the buildings as well as the positions? Describe strategies you used to try to remember the names and locations of the buildings.”

### 2.5. Experimental Procedure

Participants completed the MRI scan as part of a separate experiment either in our lab, or in another lab at the University of Pennsylvania. Then, participants were recruited to participate in the behavioral study in a separate session. In the behavioral session, participants first provided and documented informed consent, then completed the demographics and WRAT-4 measures, followed by the SAQ, SBSOD, and MRT. Then, participants completed Virtual Silcton and the debriefing questionnaire.

### 2.6. Registration and Analysis Plan

This study was registered on the Open Science Framework (OSF; https://osf.io/ea99d/) after data collection was completed for 50 participants and data analysis was completed for 33 participants. The analysis plan was created as described in the registration documents to formally establish A) one of the multiple possible ways of analyzing the data to address our hypothesis, and B) a sequential analysis data collection procedure.

We based our analysis plan on the simplest possible correlation between overall structural volume of the right hippocampus (Fischl et al., 2002) with overall pointing performance on Virtual Silcton. This analysis was chosen because it requires the least human subjectivity in data coding, and, based on the empirical literature that the right hippocampus is most likely to relate to navigation.

We proposed a sequential analysis plan because the results from 33 participants were ambiguous (a marginally significant correlation, but with a small sample size). Given the difficulty of recruiting participants with MRI data, sequential analyses allow flexibility in determining sample size. We used the extant literature to create large and small *p*-value thresholds after which data collection would stop and the results would be reported. The large *p*-value cutoff was based on the effect size between self-reported navigation ability and hippocampal volume reported by Hao and colleagues (2016) as our smallest effect size of interest. The small *p*-value cutoff was based on using *q* < .05 (*p* < .05 after applying the sequential analysis correction for multiple comparisons; Lakens 2014). We used power = 80% and would collect 20 participants batches until either we obtained a *p*-value that was smaller than *q* < .05, or until we reached 90 participants total (at which point we would have an 80% chance of detecting an effect significantly larger than our smallest effect size of interest).

Interim results that were reported on the registration on OSF used our analysis plan to determine whether additional participants would need to be recruited. Since initial registration, we learned of a pipeline that yields more accurate parcellations of hippocampal volume (which we verified with visual exploration of the hippocampal segmentations), as well as providing automated estimates of posterior and anterior volume (an exploratory question of interest). Consequently, we used this new method of anatomic analysis which was not pre-registered.

### 2.7. Statistical Tools

All processed data and code are available on the Open Science Framework (https://osf.io/ea99d/). All figures, analyses, and supplementary analyses are available in an interactive Jupyter notebook (https://mybinder.org/v2/gh/smweis/Silcton_MRI/master).

Unless otherwise specified below, statistics were calculated using the scipy and numpy packages in Python (McKinney, 2010; Oliphant, 2006). Data were manipulated with Pandas (McKinney, 2010) and visualized using Matplotlib (Hunter, 2007). Repeated measures ANOVAs were calculated using the ezANOVA package in R (version 4.4), using RStudio (RStudio Team 2016) . Effect sizes are, for *t-*tests, Cohen’s *d*, corrected for correlations for within-sample tests, and for ANOVAs, generalized eta squared (η^2^g; Bakeman 2005).

## 3. Results

We first present results from the pre-registered analyses. Next, we describe several multiple regression analyses we ran to match previous analyses (e.g., controlling for cortical volume, age, and gender). We then present exploratory analyses using the following progression. We focus on Virtual Silcton measures first, looking more broadly at subdivisions of the hippocampus (right and left) before moving on to other subcortical regions and cortical volume. Finally, we analyze non-Silcton measures, following the same progression from the hippocampus to the rest of the brain.

### 3.1. Pre-registered Analyses

Our principal analysis was the correlation between right hippocampal volume and overall pointing error on Virtual Silcton. We did not find a correlation between right hippocampal volume and pointing error, *r*(90) = .02, *p* = .88. Converting this value to a *t*-statistic yields a Bayes Factor (BF; calculated from http://pcl.missouri.edu/bf-one-sample; Rouder et al. 2009) in favor of the null hypothesis (BF_01_) of BF_01_ = 8.49. Using the original specified analysis plan (automated Freesurfer hippocampal volume calculation) to extract hippocampal volume did not change these results, *r*(90) = .07, *p* = .52, BF_01_ = 7.04.

Although we did not specify whether outliers would be excluded in our pre-registration, we re-ran this analysis excluding one outlier who had total right hippocampal volume that was approximately 4 standard deviations below the mean. Omitting this individual resulted in a slightly higher but still non-significant correlation, *r*(88) = .10, *p* = 0.38, and BF_01_ = 5.49 (see Figure 2). Using the original specified analysis plan to extract hippocampal volume did not change these results either, *r*(88) = .12, *p* = .28, BF_01_ = 4.80.

**Figure 2.**
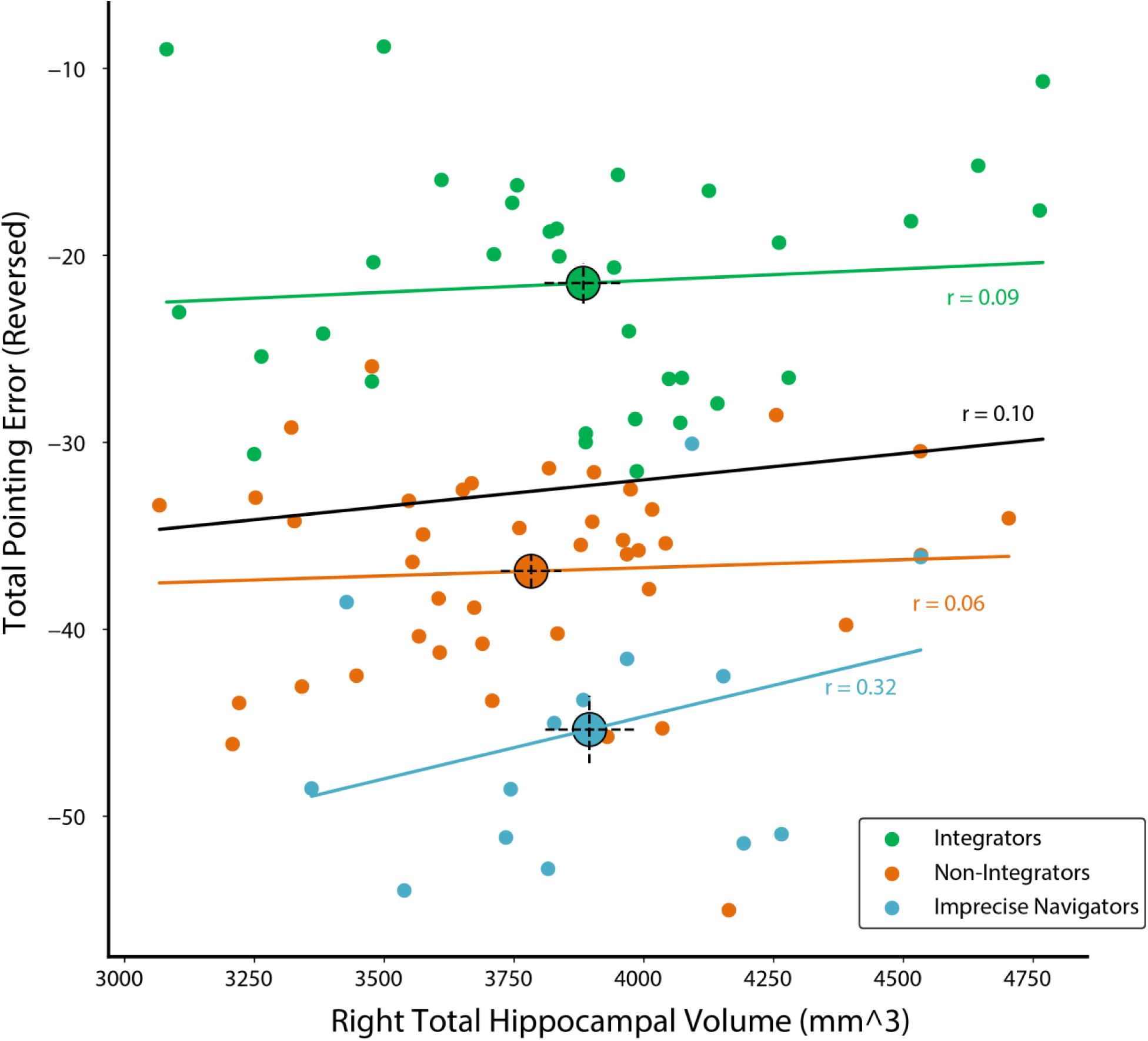
Relation between total pointing performance and right hippocampal volume. The overall correlation (black line, black font) between total pointing performance (error in degrees, reversed) and right hippocampal volume as measured by ASHS segmentation. One outlier is excluded from this scatterplot, but results were not statistically different with the outlier included. Despite numerical differences, the correlation coefficients obtained within each group do not differ statistically from each other. Large circles indicate group means and dotted lines indicate ± 1 standard error of the mean, calculated within group.

### 3.2. Multiple Regression Control Analyses

We wanted to determine whether there was a relation between right hippocampal volume and the navigation measures after accounting for cognitive and demographic factors. These control analyses are especially important because some (though not all) previous studies controlled for age, gender, and cortical volume. To account for these additional sources of nuisance variance, we ran several multiple regression analyses, controlling for gender, age, verbal IQ, small-scale spatial ability, and cortical volume. Specifically, we modeled total pointing error (and, in additional models, between-route and within-route pointing error) as a linear combination of right hippocampal (or right posterior hippocampal) volume with MRT, WRAT, gender, age, and cortical volume. No combination of regressors resulted in a significant relation between hippocampal volume and pointing error. Results of the models are reported in Table 1.

**Table 1.** Multiple regressions control analyses assessing total pointing performance with right hippocampal volume and right posterior hippocampal volume.

| Dependent variable | Predictor variable | *b* | *SE* | *t* | *p* | *R*<sup>2</sup> | Adj. *R*<sup>2</sup> |
| --- | --- | --- | --- | --- | --- | --- | --- |
| Total pointing (with total right hippocampal volume) |  |  |  |  |  | 0.21 | 0.15 |
|  | (constant) | -0.09 | 0.14 | -0.64 | 0.52 |  |  |
|  | Gender (male = 1) | 0.28 | 0.26 | 1.08 | 0.29 |  |  |
|  | Right Total Hippocampal Volume | -0.05 | 0.12 | -0.40 | 0.69 |  |  |
|  | MRT | 0.30 | 0.11 | 2.83 | < .01 |  |  |
|  | WRAT | 0.22 | 0.11 | 2.09 | 0.04 |  |  |
|  | Age | 0.15 | 0.10 | 1.44 | 0.15 |  |  |
|  | Brain Volume | 0.01 | 0.10 | 0.07 | 0.95 |  |  |
| Total pointing (with total right posterior hippocampal volume) |  |  |  |  |  | 0.21 | 0.15 |
|  | (constant) | -0.09 | 0.14 | -0.64 | 0.52 |  |  |
|  | Gender (male = 1) | 0.27 | 0.25 | 1.07 | 0.29 |  |  |
|  | Right Posterior Hippocampal Volume | -0.05 | 0.11 | -0.46 | 0.65 |  |  |
|  | MRT | 0.30 | 0.11 | 2.90 | 0.006 |  |  |
|  | WRAT | 0.22 | 0.11 | 2.08 | 0.04 |  |  |
|  | Age | 0.14 | 0.10 | 1.43 | 0.16 |  |  |
|  | Brain Volume | 0.01 | 0.13 | 0.04 | 0.97 |  |  |

*Note.* Results of two separate multiple regression analyses revealing no effect of right hippocampal volume, controlling for various other measures. SE = Standard error. MRT = Mental Rotation Test. WRAT = Wide-ranging achievement test.

### 3.3. Exploratory Analyses

We conducted several exploratory analyses. We report uncorrected p-values, but interpret findings from these analyses as exploratory results that invite replication in independent data. For interpretability, with a sample size of n = 90, a Pearson’s correlation of *r* = .21 would have a probability of *p* < .05, uncorrected. Correcting for all possible pairwise correlations (between major variables of interest) yielded a significance threshold of approximately *r* = .38. Because these analyses were exploratory, we only describe correlations as significant if they passed the Bonferroni-corrected threshold.

For hippocampal volume, we only used the results from the automated segmentation pipeline (ASHS). The remainder of cortical volume calculations come from the Freesurfer parcellation. See Supplemental Figure 1 for additional analyses using the Freesurfer and by-hand segmentation.

Similar to previous research with Virtual Silcton, we observed differences in performance on within-route pointing trials compared to between-route pointing trials (see Supplemental Figure 2). On within-pointing trials, participants pointed to a building that was on the same main route as the building they were standing near. On between-pointing trials, participants pointed to a building that was on the other main route as the building they were standing near. We analyze pointing data continuously, correlating pointing performance with brain and behavioral measures. For the correlational analyses, we collapse across between-route and within-route trials (total pointing error), but also analyzed them separately, as both show individual differences. We also analyze pointing data categorically, splitting participants into three groups – Integrators, who performed well on between-route and within-route pointing; Non-Integrators, who performed well on within-route pointing but could not integrate the two routes, performing poorly on between-route pointing; and Imprecise Navigators, who performed poorly on both types of pointing trials. We created these three groups on the basis of a K-means cluster analysis constrained to three groups on a large sample of approximately 300 participants from previous Virtual Silcton studies (Weisberg & Newcombe, 2016). Using the cutoff values from these groups yielded 34 Integrators, 42 Non-Integrators, and 14 Imprecise Navigators.

#### 3.3.1. Left and right, anterior and posterior hippocampus

Correlations between left and right total, posterior, and anterior hippocampal volumes were not related significantly with overall pointing, between-route pointing, or within-route pointing (see Figure 3). The three pointing groups did not significantly differ in total left hippocampal volume, *F*(2,87) = 0.42, *p* = .66, η^2^g = .01, nor in total right hippocampal volume, *F*(2,87) = 0.15, *p* = .86, η^2^g = .003, nor in posterior left hippocampal volume, *F*(2,87) = 2.53, *p* = .09, η^2^g = .05, nor in posterior right hippocampal volume, *F*(2,87) = 0.68, *p* = .51, η^2^g = .02, nor in anterior left hippocampal volume, *F*(2,87) = 0.05, *p* = .95, η^2^g =.001, nor in anterior right hippocampal volume, *F*(2,87) = 0.06, *p* = .94, η^2^g = .001. We also assessed the correlation between the pointing measures and the ratio of posterior to anterior hippocampal volume on the right and left. Some research has shown a link between a relatively larger posterior hippocampus and performance on navigation ability tasks (Poppenk et al., 2013). However, we did not observe such a relation (maximum *r* = -.13, correlation between left posterior-anterior ratio with between-route pointing).

**Figure 3.**
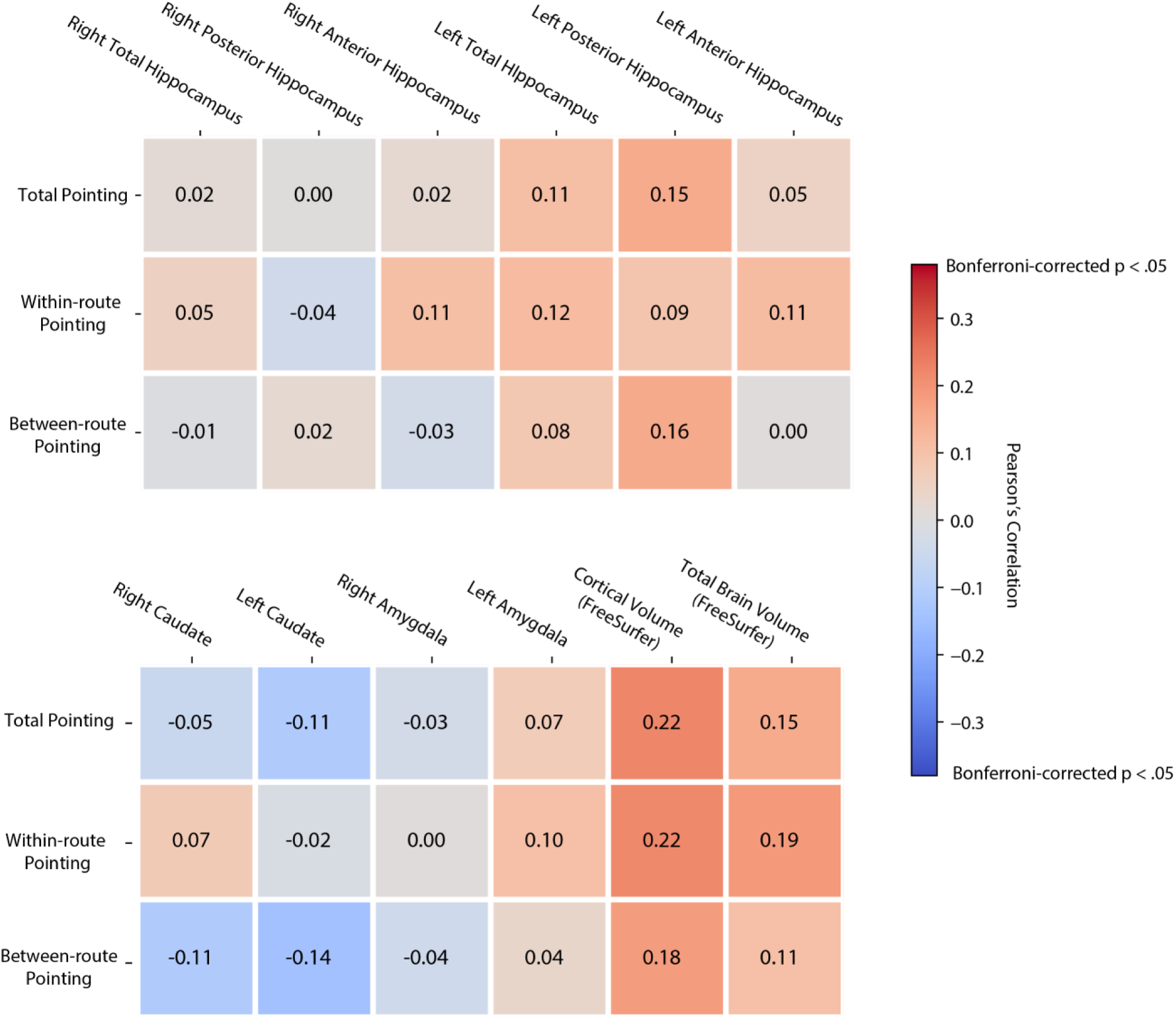
Virtual Silcton pointing correlations with brain volume measures. Pearson’s *r* correlations between Virtual Silcton pointing measures and brain measures. Hippocampal measures were calculated using ASHS, whereas additional brain area volume measures were calculated using Freesurfer.

#### 3.3.2. Cortical volume, brain volume, and other brain areas

The only notable brain-behavior correlation we observed on the pointing task was a positive relation with cortical volume, *r*(90) = .22, *p* = .037, though this did not exceed the Bonferroni-corrected threshold of *r* = .38. The caudate, amygdala, and the other medial temporal lobe structures (BA35, BA36, entorhinal cortex (ERC) or parahippocampal cortex (PHC)) resulted in non-significant correlations.

#### 3.3.3. Other Silcton measures

The model building task (measuring overall configuration or measuring within-route configuration separately) correlated negatively with hippocampal measures (-.20 < *r* < .00), despite being positively correlated with pointing performance, *r*(90) = .58, *p* < .00001. Overall, the correlation between pointing and hippocampal measures appeared stronger than the correlation between model-building and the hippocampal measures. To test this possibility statistically, we compared the correlation between pointing and each subdivision of the hippocampus (left/right, anterior/posterior/both) with the correlation between model-building and each subdivision of the hippocampus. Three subdivisions of the hippocampus were significantly more correlated (p < .05, uncorrected) with pointing compared to model-building: right anterior hippocampal volume, t(87) = 2.09, p = .04, left anterior hippocampus, t(87) = 2.65, p = .01, and left total hippocampus, t(87) = 2.53, p = .01. Right total hippocampus correlations between total pointing and model building were marginally significantly different, t(87) = 1.87, p = .06. Due to the weak correlations overall, we interpret this pattern conservatively as showing a possible differentiation of hippocampal volume as it relates to distinct types of navigational representations.

#### 3.3.4. Non-Silcton measures

We observed non-significant correlations (i.e., none surviving Bonferroni correction) between the non-Silcton measures and the volume of various brain regions (see Figures 4 and 5). Self-report measures of navigation ability, the SBSOD and SAQ, were weakly correlated with hippocampal volumes (Figure 5; -.06 < *r* < .13), whereas they were more strongly correlated with caudate, amygdala, and total cortical and brain volume (.08 < *r* < 25; see Figure 4). WRAT-4, the measure of verbal ability, was most strongly correlated with cortical volume, *r*(90) = .16, *p* = .13, and brain volume, *r*(90), *r* = .15, *p* = .16, though neither of these reached significance. Finally, the measure of small-scale spatial ability, MRT, was most strongly correlated with cortical volume, *r*(90) = .30, *p* = .004.

**Figure 4.**
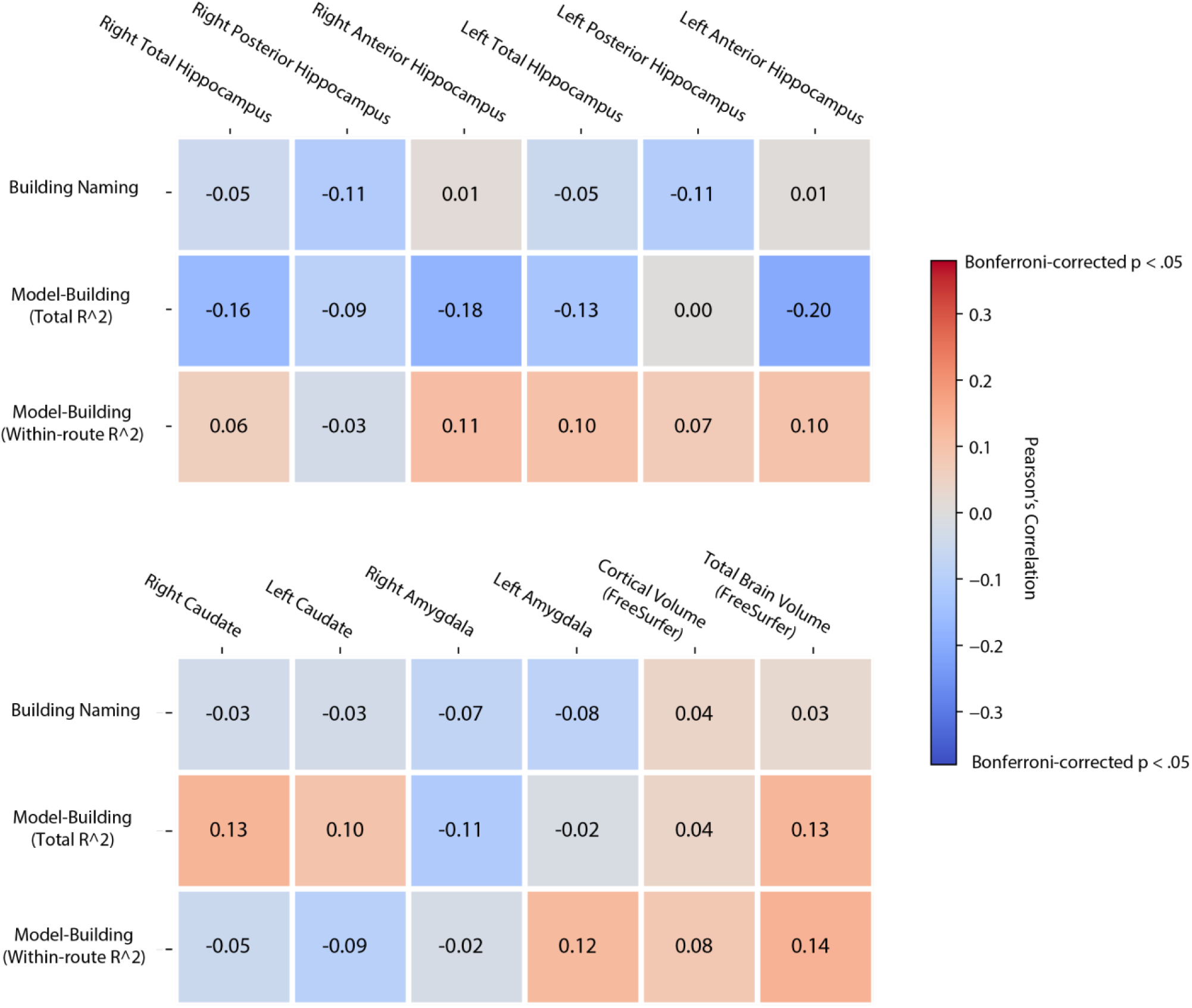
Virtual Silcton additional tasks correlations with brain volume measures. Pearson’s *r* correlations between Virtual Silcton measures and brain measures. Hippocampal measures were calculated using ASHS, whereas additional brain area volume measures were calculated using Freesurfer.

**Figure 5.**
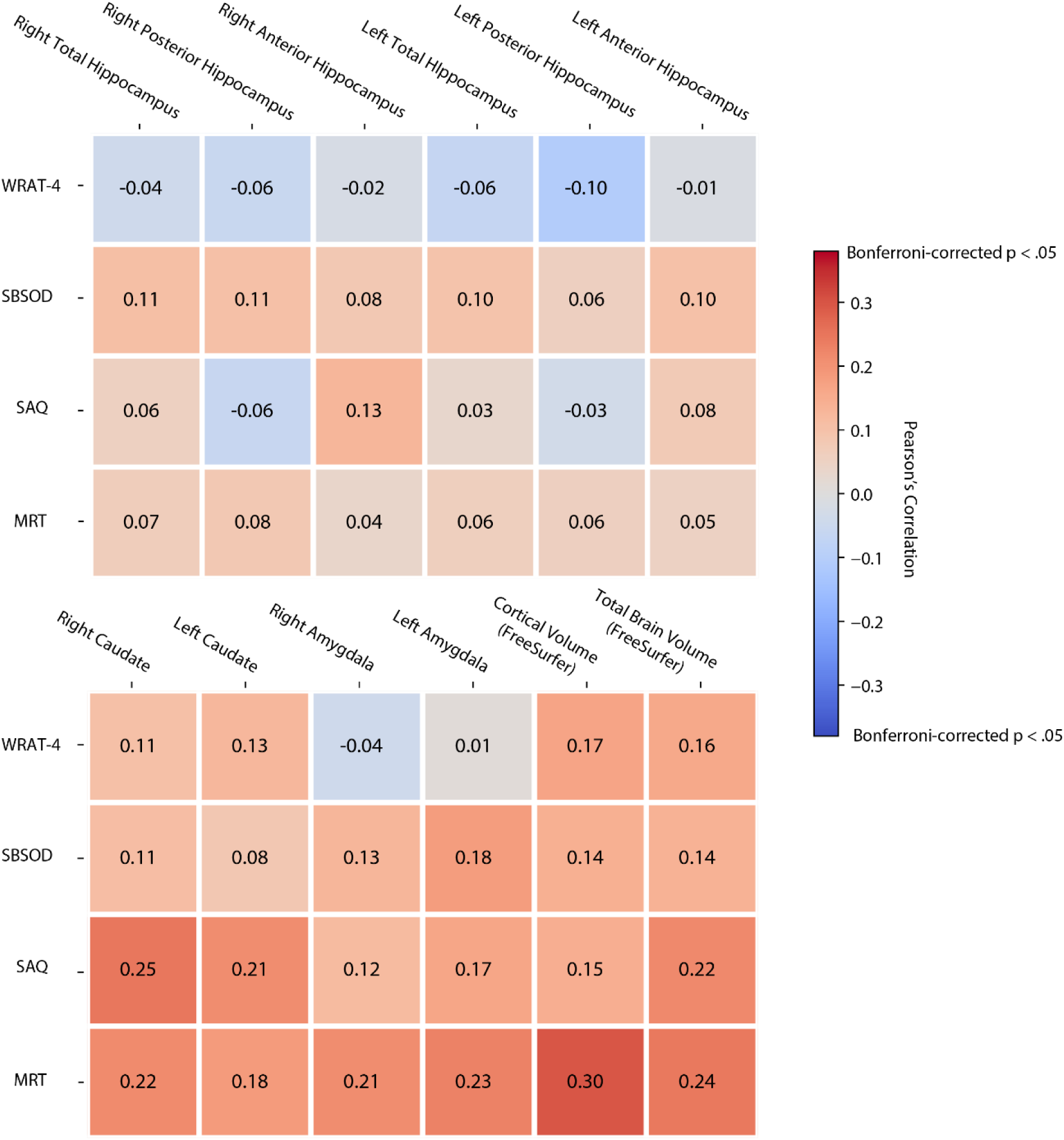
Non-Virtual Silcton tasks correlations with brain volume measures. Pearson’s *r* correlations between behavioral measures and brain measures. Hippocampal measures were calculated using ASHS, whereas additional brain area volume measures were calculated using Freesurfer. WRAT-4 = Wide ranging achievement test. SBSOD = Santa Barbara Sense of Direction scale. SAQ = Spatial anxiety questionnaire. MRT = mental rotation test.

## 4. Discussion

The hippocampus plays a crucial role in spatial navigation in humans, but the volume of the hippocampus may not be a biological marker for navigation ability among typical populations. Using an established measure of individual differences in spatial navigation we did not observe a correlation between gross anatomical properties of the hippocampus and pointing and model-building measures – two major indicators of navigation accuracy. We note several strengths of the current design. First, we used a navigational task that exhibits a wide-range of individual differences in a relatively under-studied population (in this area) of young, healthy adults. Second, we used a sample size large enough to detect small effect sizes and did so using a pre-registered analytic plan.

While it is always difficult to determine the reason for a null result, we see three possible interpretations for our results: 1) Hippocampal volume correlates with navigational ability in extreme groups, but not in typical populations. 2) Structural properties of the hippocampus and navigation behavior have a complex relationship. 3) Hippocampal volume correlates with specific skills, not general navigation ability; successful navigation also requires cognitive capabilities whose neuronal bases lie beyond the hippocampus.

### 4.1. Hippocampal volume correlates with navigational ability in extreme groups, but not in typical populations

Data from multiple sources supports the idea that expert navigators have enlarged hippocampi, whereas impaired navigators have smaller hippocampi. In humans, evidence for a link between hippocampal volume and spatial navigation ability in experts first came from studies of taxi drivers in London (e.g., Maguire et al. 2000) and in impaired navigators from individuals with hippocampal lesions (Smith & Milner, 1981). Since then, additional research by Maguire and colleagues has replicated and refined the evidence in taxi drivers, with several studies showing enlarged right posterior hippocampi relative to different control groups (Maguire et al., 2003; Woollett & Maguire, 2011), although this work suffers from small sample sizes and a non-significant interaction between right posterior hippocampal volume and right anterior hippocampal volume in taxi drivers and control groups. The association between impaired navigators and smaller hippocampal volume has also been supported in studies on pathology (Michel Habib & Sirigu, 1987; Mullally & Maguire, 2011; Packard & McGaugh, 1996), and in mild cognitive impairment and Alzheimer’s disease, which particularly affect the hippocampus and its connections (Deipolyi et al., 2007; Nedelska et al., 2012; Parizkova et al., 2018).

In the present study, we investigated navigation ability in a typical population of young, healthy individuals. Our finding is consistent with more general assessments of navigation ability, like those from self-report, that find a weak relation between hippocampal volume and navigation ability in typical populations (Hao et al., 2016; Wegman et al., 2014). One way to reconcile data from extreme groups with data from typical populations is to propose a nonlinear relation between hippocampal volume and navigation ability (see Figure 6). At the extreme ends, navigators who exclusively rely on hippocampal representations show growth in hippocampal volume, whereas navigators who cannot rely on the hippocampus (because it has degenerated or is gone) suffer the behavioral consequences. However, in the middle of the distribution, normal variability in navigation (which is nevertheless wide) is not accounted for by hippocampal volume. We consider other factors, which we discuss in the following two sections.

**Figure 6.**
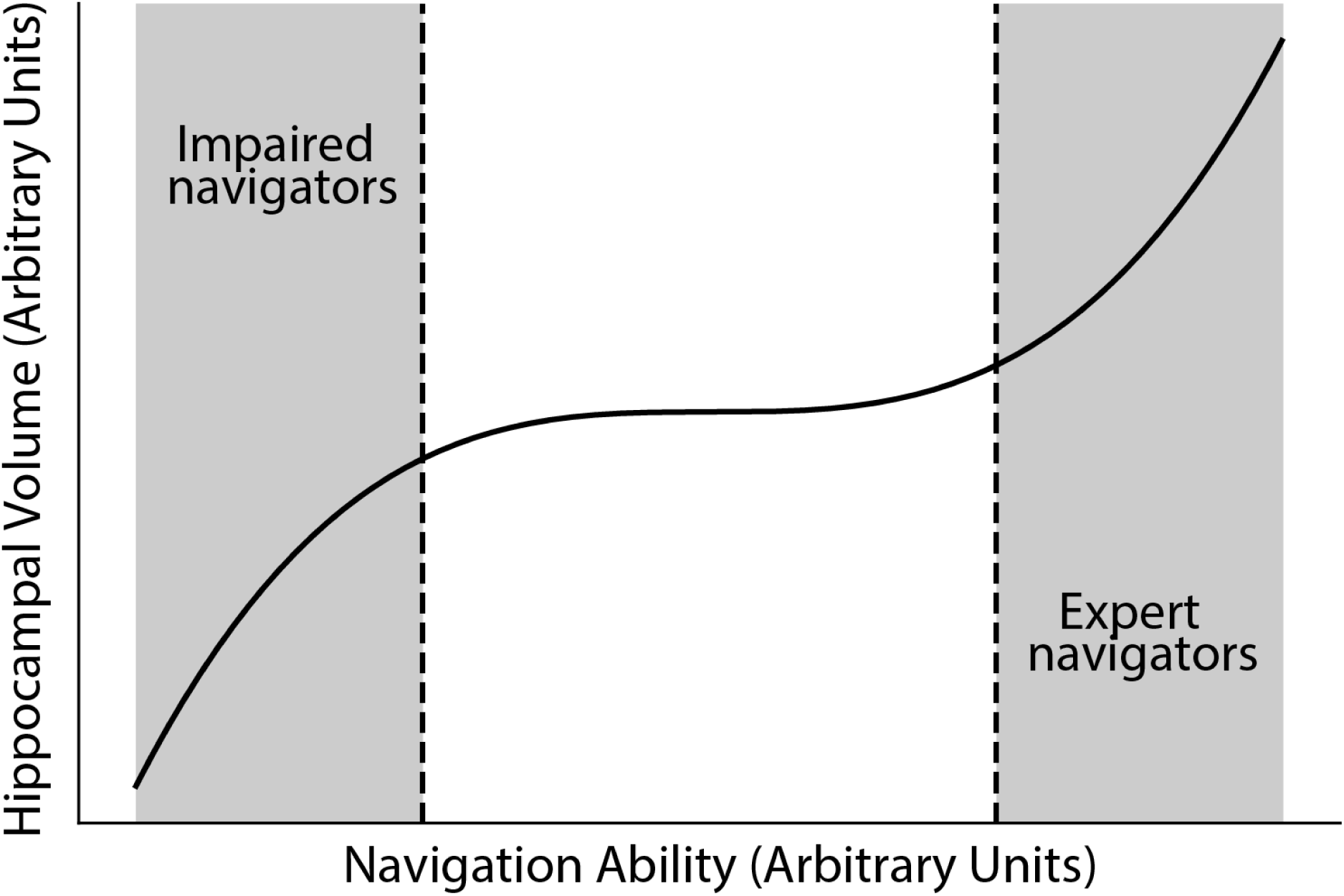
Predicted model of navigation ability and hippocampal volume across impaired, typical, and expert navigators. A visualization of the proposal that navigation ability relates to hippocampal volume in a non-linear fashion such that impaired navigators (i.e., patients with Alzheimer’s disease) and expert navigators (e.g., taxi drivers) show positive correlations with hippocampal volume and navigation ability, whereas typical populations show no or weak linear correlations. [N.B. The current study population and expert and impaired populations likely overlap on navigation ability. If the groups completely overlap, this model is implausible. If there is partial overlap, then we would expect higher correlations in expert and poor navigators, which is very weakly the case, as in Figure 2. But this assumption should be tested empirically, ideally through data collection on Virtual Silcton in patient groups and taxi drivers.]

### 4.2. Structural properties of the hippocampus and navigation behavior have a complex relationship

There are two distinct possibilities to consider here. First, the hippocampus may only be a piece of the neural puzzle, providing specific computations and processes to a network of brain regions to coordinate spatial navigation. Thus, rather than hippocampal structure relating to variance in spatial navigation behavior, changes in the connections between, for example, retrosplenial complex, the parietal lobe, and the hippocampus (Byrne, Becker, & Burgess, 2007; Ekstrom, Huffman, & Starrett, 2017; Iaria et al., 2014) may characterize information flow around the brain, which itself relates to spatial behavior. Indeed, medial temporal lobe lesions may yield greater spatial deficits when those lesions include parahippocampal cortices (Aguirre & D’Esposito, 1999; Habib & Sirigu, 1987). Similar to the difference between neural function and neural structure within the hippocampus, however, it is unclear whether individual differences in behavior would relate to connectivity structure (e.g., white matter tracts) or functional connectivity activation patterns. Given the growing literature in this area, it would be a fruitful and important avenue for future investigation.

The second possibility is that hippocampal volume may be too coarse a neuroanatomical measure to show an association with navigation ability. Aspects of navigation behavior relate only to the volume of specific subdivisions of the hippocampus, creating a complex picture of structure-function relations. Indeed, across the literature, different hippocampal properties are reported to correlate with navigation ability. Expert taxi drivers show increased right posterior hippocampal volume but decreased anterior hippocampal volume (Maguire et al., 2000) although these analyses exclude the body of the hippocampus (which we included as part of posterior hippocampus). Yet, anterior hippocampal volume correlates with path integration (Brown, Whiteman, Aselcioglu, & Stern, 2014; Chrastil, Sherrill, Aselcioglu, Hasselmo, & Stern, 2017) and self-reported navigation ability (Wegman et al., 2014). And several studies show correlations between navigation and total hippocampal volume (Hartley & Harlow, 2012; Head & Isom, 2010).

Additional data show correlations between navigation behavior and hippocampal subfields which do not follow posterior/anterior divisions (Daugherty, Bender, Yuan, & Raz, 2016). In the present study, we did not assess hippocampal subfield correlations with navigation ability because structural MRI data lacks the resolution to accurately parcellate the hippocampus into subfields (Wisse, Biessels, & Geerlings, 2014). In light of these complexities, it is unclear what if any aspects of hippocampal structure are important to navigation and what if any aspects of navigation ability relate to hippocampal structure.

### 4.3. Hippocampal volume correlates with specific skills, not general navigation ability

Perspective-taking correlates with hippocampal volume (Hartley and Harlow 2012), as does a particular navigation strategy (Konishi & Bohbot, 2013). In previous work with Virtual Silcton we observed correlations with pointing task performance and a paper-and-pencil perspective-taking test. Thus, perhaps hippocampal size is related to perspective taking, which is one (but not the only) determinant of success in navigation on Silcton. Similarly, we have found a complex relation between pointing accuracy on Silcton and navigation strategy, with integrators performing well on shortcut tasks if they choose to take the short cuts—but not all do (Weisberg & Newcombe, 2016). Again, partial overlap between hippocampal strategies and navigation accuracy on Silcton would attenuate correlations of Silcton with hippocampal volume.

Navigation ability relies on a wide range of perceptual, cognitive, and meta-cognitive processes, which likely do not all relate to the hippocampus (Ekstrom et al., 2017; Wolbers & Hegarty, 2010). In the case of the taxi drivers, it is unclear whether their expertise encompasses the creation of a cognitive map (or the capacity to do so) or maintaining an enormous catalog of associational data (e.g., recalling the names of streets, landmarks, and regions). Either of these may rely on the hippocampus, and cataloging associations correlates with the volume of hippocampal subfields (dentate gyrus and CA2/3; Palombo et al. 2018). In the case of navigation strategy, the ability to follow a familiar route through an environment, a viable alternate navigational strategy, which does not depend on the creation of a cognitive map, relies on the caudate (e.g., Marchette et al. 2011). The ability to recognize the same building from different viewpoints, which correlates with self-reported navigation ability at least, relies on representations in the parahippocampal place area (Epstein, Higgins, & Thompson-Schill, 2005), rather than the hippocampus itself.

In navigation paradigms, like Virtual Silcton, where encoding and strategy choice are unconstrained, navigation strategies that do not rely on the hippocampus could compensate for impoverished cognitive maps. Variability in these non-hippocampally mediated cognitive components of navigation could then underlie performance on Virtual Silcton in a typical population. For example, we previously showed that variation in working memory relates to performance on within-route pointing performance (Blacker, Weisberg, Newcombe, & Courtney, 2017; Weisberg & Newcombe, 2016), a process which likely does not rely on hippocampal volume or even hippocampal function.

### 4.4. Cortical Volume and Navigation Ability

Although we did not predict a correlation between hippocampal volume and cortical volume, cortical volume was the strongest correlate of the pointing task. To our knowledge, an association between navigation ability and cortical volume overall has not been reported in the hippocampal volume and navigation ability literature, although many studies correct for cortical volume when analyzing hippocampal volume. Cortical volume is associated with measures of general intelligence (Reardon et al., 2018), a finding consistent with the correlation we also observed with cortical volume and mental rotation. We emphasize that this result was exploratory, and the effect size small, but as the largest correlation we observed, we believe this effect merits further study.

### 4.5. Limitations

Several aspects of the design of the current study limit the generalizability of our results. First, although we observed a reasonable range of variability in both navigation ability and hippocampal volume, they did not correlate with each other. Nevertheless, we might speculate that the best navigators in the present sample (who arguably were at ceiling performance) would have performed worse at a more difficult task than taxi drivers; and similarly, the worst navigators in the present sample may have outperformed older adults or those with Alzheimer’s disease. Second, navigation took place in a desktop virtual reality, rather than in the real world. Although a large body of evidence supports the notion that hippocampal function can be elicited from testing in virtual environments, it is reasonable to speculate that this setting may have dampened the hippocampal contributions to navigation. Third, because we did not collect data on strategy use, nor did we have functional imaging data during navigation, we are agnostic about the strategies used by individual participants and whether their strategies engaged the hippocampus.

Future studies can address these limitations by A) collecting a more varied sample, including variations in age, general intelligence, and demographics; B) collecting data in both real-world and virtual environments; and C) collecting functional neuroimaging during the navigation and pointing phase to dissociate the role of hippocampal function from hippocampal structure.

## 5. Conclusion

In sum, this study limits the generality of the link between hippocampal volume and navigation accuracy in a typical population. These findings have implications for the role of the hippocampus in general navigation, and for the extrapolation of findings in expert and impaired groups to healthy, young adults.

## Acknowledgements

The authors wish to acknowledge Russell Epstein and Geoff Aguirre for assistance with data collection of the original MRI data.

## Abbreviations

ASHS: – Automatic Segmentation of Hippocampal Subfields
BF: – Bayes Factor
ERC: – Entorhinal cortex
MRT: – Mental rotation test
OSF: – Open Science Framework
PHC: – Parahippocampal cortex
SAQ: – Spatial anxiety questionnaire
SBSOD: – Santa Barbara sense of direction scale
WRAT-4: – Wide range achievement test 4 – verbal

